# Eyeballs on science: Impact is not just citations, but how big is readership?

**DOI:** 10.1101/136689

**Authors:** Kevin Winker

## Abstract

A full evaluation of the impact of scientific publications needs to count readers who don’t generate citations. But this is very difficult to do. I set up a scholarly foraging experiment to try to estimate readership for my own body of work: a personal web site with ‘reprint’ pdf files available for downloading and a web statistics program engaged to count these downloads. (I made the site topically rather than author-oriented to increase potential audience size.) Despite this, human users are difficult to count. There are a lot of bots, spiders, and other web programs that increasingly mimic human behavior (e.g., with IP- and chrono-camouflage), and humans’ own behaviors are changing (e.g., reading papers online many times). From four years of data, after culling both automated activity (a fascinating ecosystem) and humans reading repeatedly online, my papers receive ∼4000-8000 downloads per year, about an order of magnitude higher than the number of citations they receive annually. This is a conservative minimum, because these publications can be obtained from other sources as well. On average, this body of work may be being read at an approximately 10:1 download-to-citation ratio. At a more granular scale, downloads do not correlate with citations; there are some papers being downloaded at about a 100:1 ratio, and it turns out these are exactly how they were meant to be used (e.g., an instruction manual). How intensively someone reads a paper is of course highly variable, but this exercise gives us an idea of how broad our publications’ impacts actually are. Citations alone don’t come close to measuring it.

## Introduction

The evaluation of scientific impact has dwelt heavily on the number of times a work is cited.This is not unreasonable, and citation statistics have been readily available for most scientific journals through data aggregators such as the Science Citation Index (Thomson-Reuters), Scopus (Elsevier), and Google Scholar. But we know that this does not fully describe how often a work is read. How many of us read scientific papers that we don’t cite? Clearly, measuring our consumption of science is not fully done using citation metrics. And consider also that fellow research scientists are not our only audience. In communicating our science, a broader audience of students, instructors, and the public is also important. How frequently our works are used beyond simple citations is an important question to answer in the development of more comprehensive evaluation of impact. And a more complete evaluation of impact is useful both to authors and to their evaluators. How different is consumption from citations? And how might we measure this?

In the transition from physical to electronic distribution of scientific publications, it became possible to partially account for the number of times individual articles were accessed. This ability probably achieved its earliest transparency when journals went entirely online and openly reported the number of times an article was viewed. These metrics have been rapidly improving, giving us an increased ability to see how often articles are viewed, downloaded, cited, and discussed or used in various electronic media (often termed ‘altmetrics’ and posted prominently with an online article). As individual authors, however, these metrics are not yet widely available to us, leaving patterns and degrees of non-cited use an open question.

Even within one’s discipline, uses that don’t generate citations may be even better than those that do. For example, who wouldn’t prefer to have their work be the focus of a journal club discussion rather than garner a throwaway citation, e.g., as a reference to someone who’s recently worked on a particular topic?

Over the past five years, I ran a scholarly foraging experiment to begin to evaluate this issue for publications upon which I was an author. I didn’t have even a foggy sense of the magnitude of the readership these publications were reaching. My goals were to measure the size of this phenomenon relative to citation rates and to see what degrees of variation existed among individual papers.

## Methods

My experiment involved posting my publications on my web site and paying my hosting company to implement web count statistics (using AWStats, a commonly used open-source program; www.awstats.org) so that uses were archived in web log files. This sounds very simple. It is not.

I already had a crude, CV-style web page and my own web site because I had found that the resources my university provided at the time were unacceptable (e.g., their web statistics and reporting were not up to my standards). I recognized, however, that I wanted to accommodate *topically* oriented rather than *author*-oriented searches. A personal web page is more likely to be the latter, and that immediately narrows the universe of potential readers to (mostly) those who might know of you and your work. So I began by annotating my publications list, essentially adding brief abstracts of the papers’ contents (not identical to the published abstracts). This substantially boosted the search target size of my scholarly foraging experiment (Fig. 1). What really boosted it, though, was adding pdf files and allowing search engines to index those files. The core of the scholarly foraging experiment is thus a cafeteria experiment, with the papers themselves as the various bits of (intellectual) nutrition. This created a search target size for my body of work greatly exceeding anything similar (e.g., my Google Scholar profile, which I do not make public so as to not have acompeting, uncountable search target; Fig. 2). With this done, the experiment was open for business, both to other researchers who know of our work and to users interested in the primary literature on topics x, y, and z when those search engine parameters turned up something we’d done. This increase in search target size was an unanticipated fringe benefit of lasting value (as seen in AWStats summaries of search terms used to find the site/paper).

**Figure 1.**
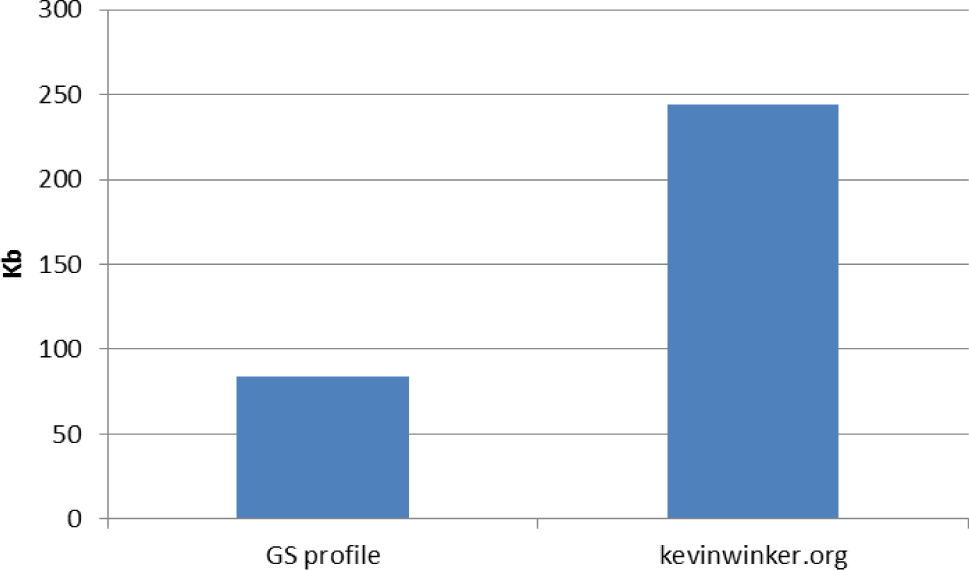
Comparing search target sizes of two sources prior to posting reprints.

**Figure 2.**
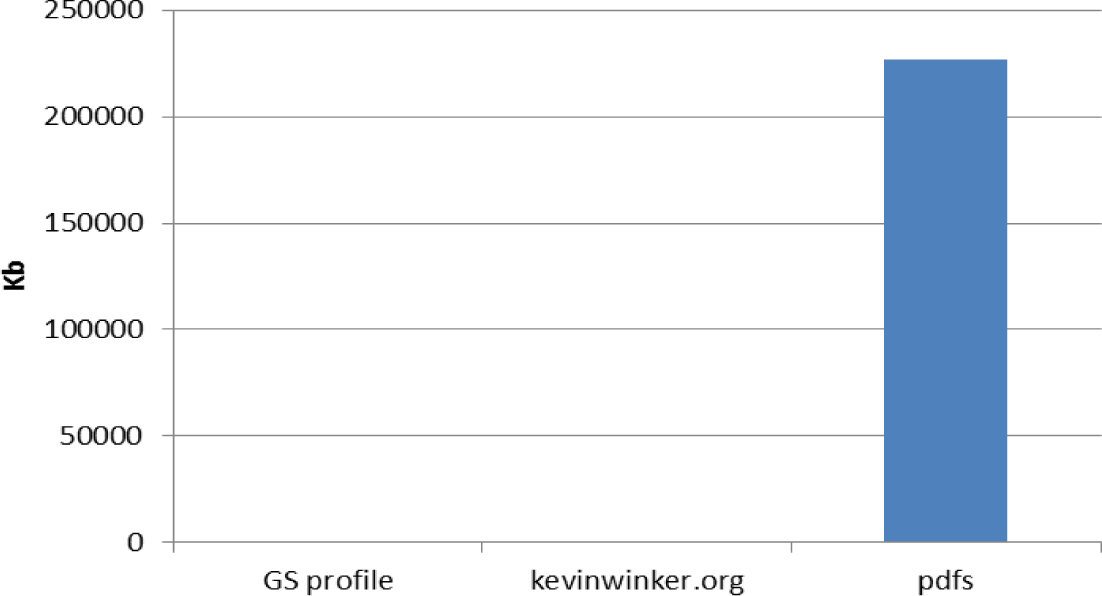
Comparing search target sizes. Adding searchable pdf files to the.org site created a very large target for search engines, effectively creating a large, topically oriented (rather than author-oriented) site.

I could immediately see from online AWStats summaries that our papers were being downloaded. The rates seemed very high, though, with a puzzling imbalance between unique visitors and downloads. Agents being counted as unique visitors were taking, on average, a seemingly high number of pdf files. It turns out that while the AWStats online interface states that it is excluding bots and spiders from its summary tables, for download statistics this is not the case:these automated activities are also included in the download summaries. So I had to ask “How many humans are downloading papers?” There was nothing I could do to obtain accurate counts from the online interface. I initially let this run for about two years, though, before digging in detail through the accumulated log files. This was the first of two such detailed engagements.

My site generates about 100,000 lines of log files each year. I went through them manually after pulling them into a spreadsheet so I could sort in multiple ways to get a feel for human versus automated activity. It is remarkably complex (more on this below and in the Appendix). As a field biologist, I saw a fascinating electronic ecosystem with a surprisingly diverse community of human and non-human actors. The best automated ones (e.g., Google, Baidu, Yandex, Yahoo, Bing, etc.) label themselves as such and are thus easily culled. But many fly unlabeled, and these need to be spotted by their behavior (e.g., downloading four pdf files in five seconds).

After doing this for days with two years of data, I decided to add something to make it easier to find the automated actors in future log files. As a field biologist, I wanted to have a component in the system that would cause automated actors to self-mark. So I put in what I called honeytraps: nice, juicy pdf files made up of totally random text that spanned the file sizes of my real pdf files. These honeytraps would have zero nutritional value for human actors but should be snatched up gleefully by bots and spiders. (I called these ‘mud’ files, because I can’t think of anything more useless to bring home from the field—not just dirt, but heavy, wet dirt.) It worked pretty well.

I’ve now gone through four of the five years of data, the first two and the last two years. It is a fascinating ecosystem to study, but it is time consuming to cull automated activity. I don’t think it would be possible to write a program that could weed out automated activity better than I can do it manually. And it turns out that this is a tough nut to crack for professionals (e.g., see Paola Kathuria’s blog entry here: www.limov.com/library/do-not-believe-your-web-stats.lml). One surprise was how much things changed over this period, both in automated and human behaviorsBoth are evolving, and both are exhibiting more traits of the other, making it increasingly difficult to distinguish between them. Programs have been using IP-address camouflage for a while (coming in under different IP addresses to avoid detection), but now some include chrono-camouflage, trying to mimic a human in frequencies of activity. And humans are increasingly using my site’s storage space rather than their own and returning multiple times to read the same paper online. A really surprising change over just three years was the sharp increase in how many humans read papers on their iPhones. I do not think getting a highly accurate count of human readership is possible. But these rather laborious methods do work to obtain reasonable, minimum-use estimates.

## Results

After culling automated activity and humans reading repeatedly online from four years of data (the first two and last two of the five-year period), my papers receive ∼4000-8000 downloads per year. This is about an order of magnitude higher than the number of citations they receive annually. This is a conservative minimum, because these publications can be obtained from other sources as well. On average, then, this body of work may be being read at an approximately 10:1 download-to-citation ratio.

Comparing downloads per paper with their citations gives a useful granular view of how variable this use ratio can be (Fig. 3). Downloads do not correlate with citations, and some papers in this body of work are being downloaded at about a 100:1 ratio (Fig. 3). In considering which these are, it turns out that this is exactly how they were meant to be used. For example, one is an instruction manual, and another is a paper most useful in study design and perhaps proposal writing.

**Figure 3.**
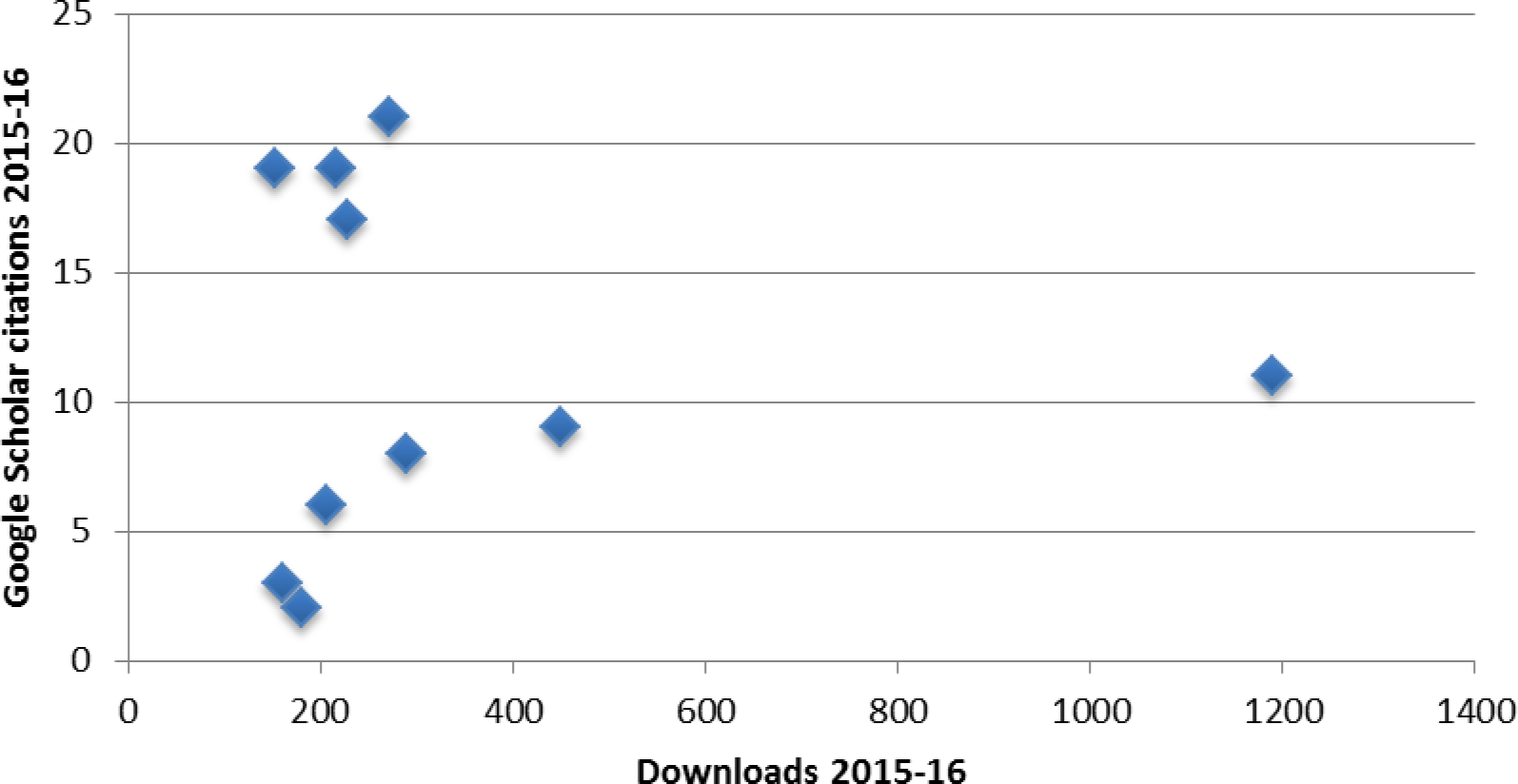
The top ten downloaded files and their Google Scholar citation rates during the same period (2015-2016). Note the lack of correlation and the variation in the download:citation ratios from ∼10:1 to∼100:1.

## Discussion

It appears that the readership for this body of work is, conservatively, on average about ten times as large as the number of citations indicate. Having an idea of the magnitude of exposure and thus possible readership is useful information. Even more useful to me as an author is that for some papers the readership can be as much as 100 times larger. It is reassuring to see the intended effect of papers written more for an audience to be educated on a topic rather than for an audience likely to generate citations on that topic. The longer-term popularity of some of these (data not shown) is also illuminating. The top ten downloads in 2015-16 had publication dates ranging from 1998 to 2010 (averaging 2007).

Despite much development in this area of non-citation quantification over the past five years (e.g., increasing adoption of altmetrics), as yet I see no substitute for such an involved personal engagement for studying the impacts of one’s own body of work. ResearchGate is useful, but it has taken a long time to grow out of being a small, gated community (readership numbers are increasing, though it is not possible to separate bots if there are any). Altmetrics are wonderful, but they don’t exist for older works, they are not used by all journals yet, and there is no aggregator to obtain them all in one place.

How intensively or carefully someone reads a paper is of course highly variable, but this exercise gives us an idea of how broad our publications’ impacts actually are. Citations alone don’t come close to measuring it.

## Appendix

If you want to try a similar experiment yourself, here are some things that I learned that might help.

robotx.txt files are not reliable to label bots, spiders, etc. Many ignore it. Nor are.gif files. There are highly specialized.gif hunters, and those tend not to pick up.pdf files (and vice-versa).

There are some really good programmers out there building bots that completely avoid my honeytrap mud files. They must have some sort of content interpretation capacity. And there was one over a five-week period in 2015 from Russia that used IP address camouflage and only consumed one single mud file, repeatedly. I called it the Russian camouflaged honeyeater and had a mental image of some Frankensteinian monster periodically released from the coding dungeons of Skynet to engage in absolutely bizarre feeding. The bots that only hunt.gif files are odd, too.

It is the multi-reading humans that are among the most difficult to accommodate. Their number increased over the course of my experiment. In the first two years I culled anyone coming back more than twice, reasoning that others were like me in recognizing when they had already seen (and perhaps saved to their storage space) a paper and gotten what they needed there. In the last two years of data there were many more clearly human users that came back repeatedly.

The changes in this ecosystem over time, among human and automated actors, make static code to accurately count humans virtually impossible to create from my perspective. Keeping up dynamic code to do so would require a great deal of time and would likely possess considerable commercial value. So we are unlikely to ever get accurate numbers of human readers. From a single download for a journal club group to the difficulty of counting electronic users (e.g., one lab computer being used by many students, or one person using multiple computers or a having a dynamic IP address), there are not really reliable ways to count uses, so we’ll probably always have only approximations.

